# Childhood maltreatment and brain aging during adulthood

**DOI:** 10.1101/2025.01.16.633271

**Authors:** Leland L. Fleming, Kyoko Ohashi, Michelle Bosquet Enlow, Jennifer Khoury, Torsten Klengel, Karlen Lyons-Ruth, Martin Teicher, Kerry J. Ressler

## Abstract

**Importance:** Childhood maltreatment (CM) is associated with the early onset of psychiatric and medical disorders and accelerated biological aging.

**Objective:** To identify types of maltreatment and developmental sensitive periods that are associated with accelerated adult brain aging.

**Design:** Participants were mothers of infants recruited from the community into a study assessing the effects of CM on maternal behavior, infant attachment, and maternal and infant neurobiology. Data were collected from July 2015 to November 2019 and were analyzed from July 2023 to October 2024.

**Setting:** Academic medical centers.

**Participants:** High-quality MRI scans were obtained on 92 of 150 mothers enrolled in the study. The main exclusion criteria for neuroimaging were histories of head trauma with loss of consciousness or concussion, psychotropic use before age 18, pregnancy, and customary MRI exclusions (e.g., metal implant). The primary reasons for non-completion of the neuroimaging study were unwillingness to be scanned, inability to attend the MRI study visit due to work and/or childcare, metal implants, or pregnancy.

**Main Outcome(s) and Measure(s):** The Maltreatment and Abuse Chronology of Exposure scale was used to retrospectively assess the annual severity of exposure to ten types of CM from birth to age 18 years. Brain age was calculated from T1-weighted 3T MRI Scans using a previously published machine learning algorithm. Sensitive periods were identified using random forest regression with conditional inference trees.

**Results:** Forty-nine (53.3%) of the 92 mothers (mean [SD] age, 32.4 [4.3] years) reported experiencing one or more types of CM. Total CM severity was associated with accelerated brain aging (β=0.05, 95% CI, 0.02 to 0.09, p<.005). The most robust type/time risk factors for accelerated brain aging were parental physical abuse between ages 4 to 6 years, witnessing sibling violence between ages 4 to 15 years, parental verbal abuse between ages 10 to 12 years, and parental emotional neglect between ages 16 to 18 years.

**Conclusions and Relevance:** Several types of CM between ages 4-18 years were associated with accelerated brain aging. Understanding how these specific types and ages of exposure contribute to accelerated brain aging may provide important insights into preventing key clinical consequences of CM.

**SUMMARY:** **Question:** How does childhood adversity relate to brain aging in adulthood, and are there sensitive periods for this association?

**Findings:** In this cohort-based study of adult women, we observed sensitive periods for the association between adult brain aging and five classes of childhood maltreatment: parental physical abuse, parental verbal abuse, parental emotional neglect, and witnessing sibling violence.

**Meaning:** These findings suggest that maltreatment subtype and age at exposure may be important factors contributing to the impact of childhood adversity on brain aging later in life.

## INTRODUCTION

Early life experiences significantly impact nervous system structure, development, and function, leading to profound impacts on health and disease ^1–3^. Negative experiences in particular, like childhood maltreatment, can have severely detrimental and chronically persistent consequences ^3,4^. The US Centers for Disease Control and Prevention (CDC) have reported that childhood maltreatment accounts for a large proportion of the risk for drug addiction (64%) ^5^, depression (54%) ^6^, and suicide attempts (67%) ^6^. Moreover, childhood maltreatment is associated with a reduction in life span of as great as 20 years ^7^. These markedly adverse outcomes in psychiatric health and mortality have created a need to prevent and reverse the detrimental effects of childhood maltreatment. However, a fundamental understanding of the factors contributing to differential outcomes among maltreated individuals is still needed.

One long-term effect of early life adversity is an overall acceleration in development and aging ^8,9^. This association is thought to be due to increased allostatic load, i.e., the aggregated wear and tear of adverse experiences on the body ^10^. Biological aging measures, such as “brain age” ^11,12^, allow for the quantification of these cumulative effects. Brain age uses age-associated structural features across the brain to derive the apparent “age” of the brain. The difference between an individual’s brain age and chronological age, referred to as brain age gap (BAG), predicts mortality, with accelerated brain aging associated with increased mortality risk ^12^.

Previous work suggests that pediatric brain aging is also linked to experiences of early life adversity ^13–15^. However, to the best of our knowledge, no published studies to date have examined the relationship between childhood maltreatment and brain aging in adulthood.

Evidence suggests that the age at which childhood maltreatment is experienced may significantly influence its effects on mental health symptoms ^16,17,18^. This phenomenon mirrors other developmental processes with “sensitive periods,” during which the brain’s susceptibility to environmental influences is heightened^19^. Current literature suggests that the timing of maltreatment exposure is particularly important when examining its effects on neurobiology^20–25^. For example, sensitive periods for the effects of maltreatment have been identified for emotional circuitry regions in the brain. The hippocampus shows peak sensitivity to abuse during early life (∼3-7 years), with a second peak during adolescence (10-16 years)^21,25^. Increased amygdala volume is associated with childhood maltreatment during preadolescence (10-11 years) ^22^.

Sensitive periods for the function of these structures have also been identified, with hyperactive responses to threatening stimuli associated with maltreatment exposure during the teenage years, and hypoactive responses linked to exposure during early childhood^20^. Still, it remains unclear how childhood maltreatment, and specifically the timing of exposure, impacts cumulative measures of brain development, such as brain age. Moreover, the long-term impact of childhood maltreatment on brain aging into adulthood is not known.

In the present study, we aimed to investigate the association between childhood maltreatment and brain aging in a sample of adult women with infants. We additionally examined whether any observed associations appeared to be influenced by sensitive periods of exposure. In doing so, we aimed to better understand the relations between childhood maltreatment and brain development while also expanding our knowledge of how different windows of neuroplasticity may influence later outcomes associated with early life adversity.

## METHODS

### Study participants

Participants were drawn from the Mother-Infant Neurobiological Development (MIND) study, designed to assess the impact of maternal childhood maltreatment on maternal and infant neurobiological and behavioral outcomes. Of the 150 eligible mothers in the MIND study, 19 were unresponsive to follow-up efforts via phone and email communication, 18 declined MRI participation, typically because of work schedule or childcare responsibilities, and 5 were disqualified based on MRI exclusion criteria. Hence, 102 participants were invited for MRI. Three missed appointments without rescheduling, and 3 experienced complications (panic attack, nausea, claustrophobia) that precluded scanning. Of the 96 participants who completed the scan session, 4 scans were unusable due to excessive motion or imaging artifacts. High-quality scans were obtained on 92 mothers (mean age = 32.38 (+/- 4.27) years, years of education 17.80 = (+/- 3.16) years, racial breakdown: 7 Asian American, 8 Bi/Multi-racial, 6 African American, 70 Caucasian, 1 Native Hawaiian/Pacific Islander). All study procedures were carried out under the Institutional Review Board of the McLean/Mass General Brigham Healthcare system (IRB Protocol #: 2014P002522). (See supplemental methods, table S1 for additional details).

### Maltreatment history assessment

The Maltreatment and Abuse Chronology of Exposure (MACE) Scale ^26^, was used to retrospectively assess the timing and severity of maltreatment. The MACE-X consists of 75 items, and participants were instructed to indicate whether they experienced a given event and, if so, to indicate the ages at which the event occurred. Rasch-based severity scores (ranging from 0 – 10) were provided for ten types of maltreatment across age, including parental physical abuse, parental verbal abuse, nonverbal emotional abuse, sexual abuse, emotional neglect, physical neglect, witnessing interparental violence, witnessing violence towards siblings, peer emotional abuse, and peer physical bullying. Scores were also provided for each maltreatment category based on the items endorsed regardless of age, as well as an overall composite severity score that summed each of the maltreatment types (range 0 – 100). The MACE has excellent test-retest reliability across all ages and each maltreatment subtype (r=0.91) ^26^ and has been used to demonstrate sensitive periods for associations of maltreatment exposure with psychopathology^27–29^ and with brain structure ^22,25^ and function ^20,30^.

### MRI data collection

Neuroimaging was performed on a Siemens Magnetom Prisma (3T; Siemens AG, Siemens Medical Solutions) using a 64-element phased-array radiofrequency reception coil. A high-resolution 3D T1-weighted sequence was acquired for anatomical registration (TR = 2530 ms, TEs = 1.69–7.27 ms, flip angle = 7°, TI = 1100 ms, voxel size = 1.0 × 1.0 × 1.0 mm).

### Brain age calculation

Brain-predicted age was calculated using a previously published machine learning model ^12,31,31^. The method utilizes a model based on Gaussian process regression to predict brain age using structural features from T1-weighted MRI images. The algorithm was originally trained on 3,377 healthy individuals (ages 18-92 years, mean age = 40.6 years, SD = 21.4), and tested on 857 individuals (18-90 years of age, mean age = 40.1 years, SD = 21). Following standard pre-processing (details in supplement), we quantified brain aging by determining the brain age gap (BAG), or the disparity between brain-predicted age and chronological age (see details in *Statistical Analysis* section).

### Brain age model validation

Following the calculation of brain age, we confirmed the overall prediction accuracy of the model in our dataset by calculating the mean absolute error. This was done to ensure that the brain age model, which was trained in a different dataset (described above), could be accurately applied to our specific sample of participants. Here, we obtained mean absolute error values comparable to other larger datasets, including the UK Biobank ^32^ and data originally used in the validation of the brain age model ^11^ (see Results section).

### Statistical analysis

To calculate brain age gap (BAG) we took the residual values from regressing brain-predicted age on chronological age in line with similar studies of biological aging ^32,33^. This approach allows for examining residual variations in brain age independent of chronological age and is an important adjustment necessary to avoid the possible overestimation of age in younger brains and an underestimation of age in older brains ^33–35^. We then compared BAG to their composite severity score using a negative binomial regression model, given its ability to robustly handle potentially zero-inflated distributions, as in the case of maltreatment data, as well as to mitigate issues of overdispersion (variance greater than the mean) ^36^. Follow-up sensitivity analyses were conducted by adding covariates (household income, age, total cortical volume, and smoking status) based on their previously documented associations with brain aging ^37–39^.

A random forest regression with conditional inference trees was conducted to determine whether severity of exposure to specific types of childhood maltreatment at specific ages is associated with accelerated brain aging in adulthood. This technique was chosen because it does not assume a linear relation between the response and predictor variables ^40^, can handle larger numbers of predictor variables with high levels of collinearity, and has higher performance compared to similar approaches (e.g., support vector machines, neural networks, and gradient boosted machines) in accurately identifying the most important predictors of neuropsychiatric outcomes ^25,29,41^. More details on this method are provided in the *Supplemental Methods*.

In each model, for each maltreatment subtype, the severity score and age of occurrence were used to predict BAG with covariates of household income, chronological age, total cortical volume, and smoking status included, based on previous literature ^37–39^. Developmental windows of 3-year increments were used to reduce data dimensionality, similar to other work ^16,42^.

Multiple comparisons correction was applied by calculating Bonferroni adjusted p values. To determine directionality, we used the saved random forests and modified the exposure for each predictor variable from zero to maximum while holding all other predictors constant at their modal value in the model ^25,30^. Subsequent negative binomial regressions were also run to determine incident rate ratios (IRRs) for the severity of exposure to each maltreatment subtype (Table 2). Note: IRR values were generated to more clearly show percent changes, but statistical significance is based solely only upon the results of the random forest models.

## RESULTS

### Brain age acceleration and cumulative childhood maltreatment

We first assessed the general accuracy of the brain age metric by calculating the mean absolute error (MAE) of the brain age model in our dataset (Fig. 1A). We obtained MAE values (4.68 +/-3.37 years) comparable to those obtained in the analysis of other datasets, including the UK Biobank ^43^ (MAE range: 4.6 to 5.5 years) and data originally used in the validation of the brain age algorithm ^12^ (MAE = 5.02 years). Additionally, brain-predicted age showed a significant correlation with chronological age (R=0.52, p=8.93e-08). Brain age gap (BAG) was quantified by taking the residuals from modeling brain age on chronological age as described in the Methods section (Fig. 1B). We then assessed the association between total childhood maltreatment exposure severity and BAG (Fig. 1C). We observed an overall positive association between the composite severity score and BAG (*β=.05,* p<.01). A plot of the association between BAG and total childhood maltreatment is shown in Fig. 1C. A follow-up sensitivity analysis was then performed to test whether the results were maintained after controlling for covariates. In these analyses, the main results held, with the overall severity of childhood maltreatment still associated with greater BAG (*β = .05,* p<.05).

**Figure 1.**
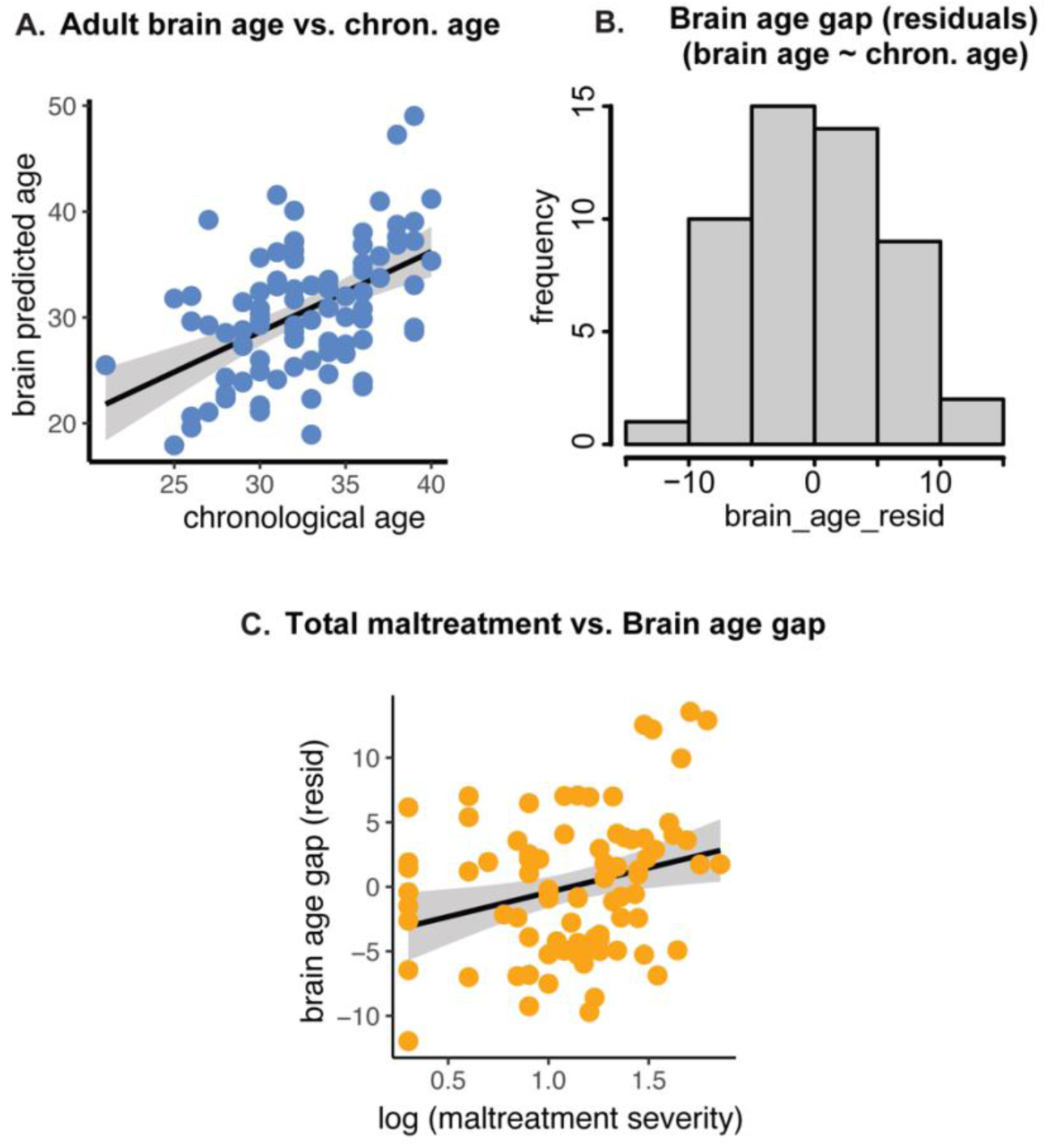
Childhood trauma and brain age acceleration in adulthood. The accuracy of the brain age metric by calculating the mean absolute error (MAE) of the brain age r model in our dataset (panel A). We obtained good model fit as demonstrated by low MAE values (4.68 +A3.37 years) and a significant correlation between brain and chronological age (Pearson’s R=0.52, p=8.93e-08). The distribution of the brain age gap values is shown as histogram in panel B. The association between composite maltreatment severity and brain age was also assessed and is shown in panel C. The regression model revealed that greater maltreatment severity was significantly associated with accelerated brain aging (β = .05, p=3.48e-02) after controlling for covariates of income, age, total cortical volume, and smoking status. For visual­ization purposes, fog transformed maltreatment severity is shown.

### Sensitive periods for effects of maltreatment on brain aging

Next, we tested for the presence of sensitive periods during which the effects of childhood maltreatment on brain aging were most prominent. We implemented a random forest regression model with conditional inference trees to detect the effects of age of exposure and maltreatment subtype on BAG. This model included covariates of household income, total cortical volume, and smoking status. Overall, four subtypes of maltreatment exhibited significant peaks suggesting timing sensitivity: parental verbal abuse, parental physical abuse, witnessing sibling violence, and emotional neglect (Fig. 2, Table 1, supplemental table S2). A sensitive period for parental physical abuse was detected in the 4- to 6-year age range. Witnessing violence towards siblings showed a wide range, with the sensitive period spanning 4 to 15 years of age. Parental verbal abuse exhibited a sensitive period from ages 7 to 12 years. A sensitive period for emotional neglect was also observed, with a window from 16 to 18 years.

**Figure 2.**
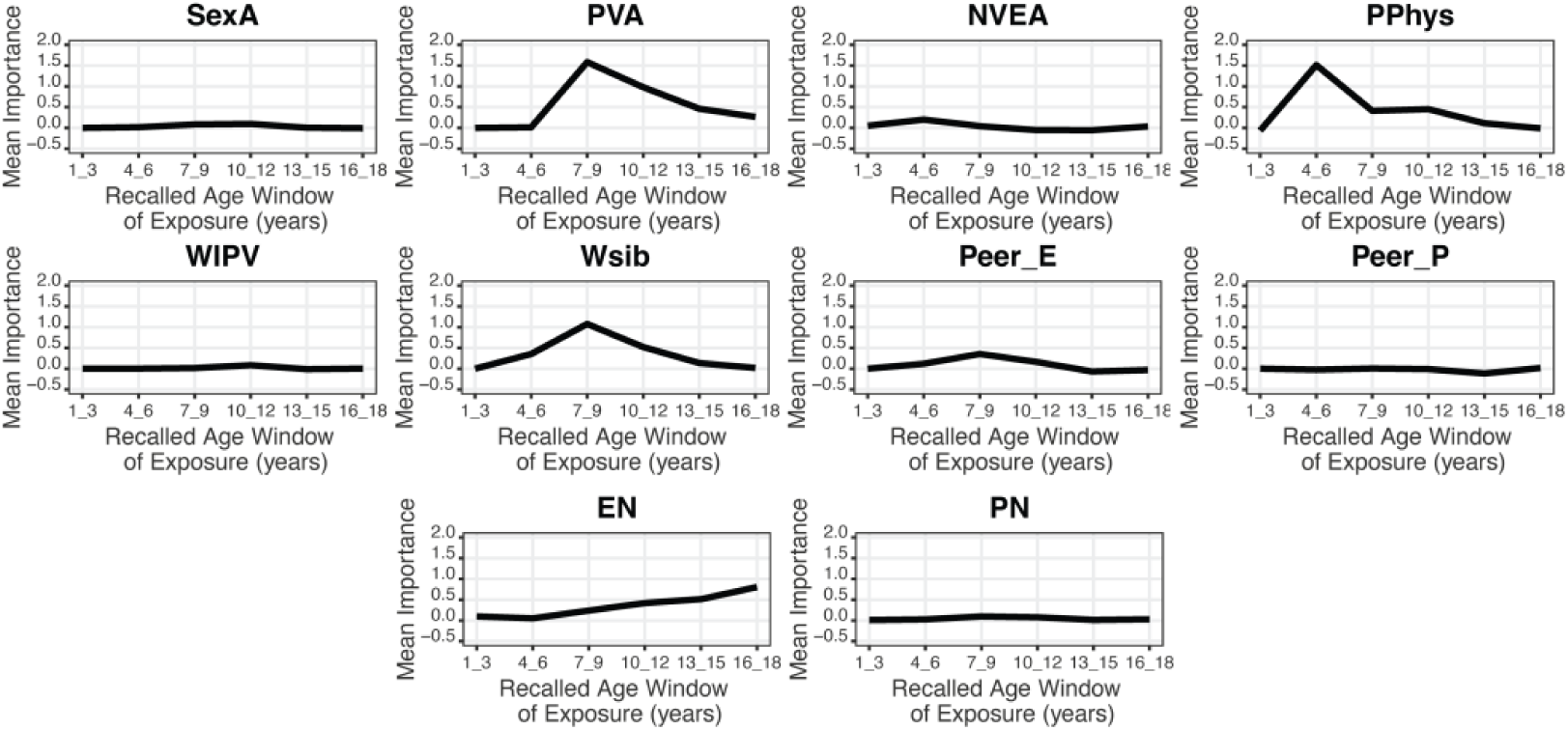
Sensitive periods by maltreatment subtypes. A random forest regression with conditional inference trees was conducted to gain a more detailed view into how exposure to specific subtypes of maltreatment influences brain aging during adulthood. The horizontal axis for each plot shows the recalled age window of exposure to each maltreatment in years. Vertical axes show mean importance, defined as the average rise in the mean square error of the model’s fit after permutation of each individual variable. The ten maltreatment subtypes examined from the MACE questionnaire are displayed. Abbreviations are as follows: PVA, parental verbal abuse; NVEA, nonverbal emotional abuse; PPhys, parental physical abuse; WIPV, witnessing iπterpa-rental violence; and WSIB, witnessing violence toward siblings; Peer E, peer emotional bullying; Peer_P, peer physical bullying; EN emotional neglect; PN, physical neglect.

**Table 1.**
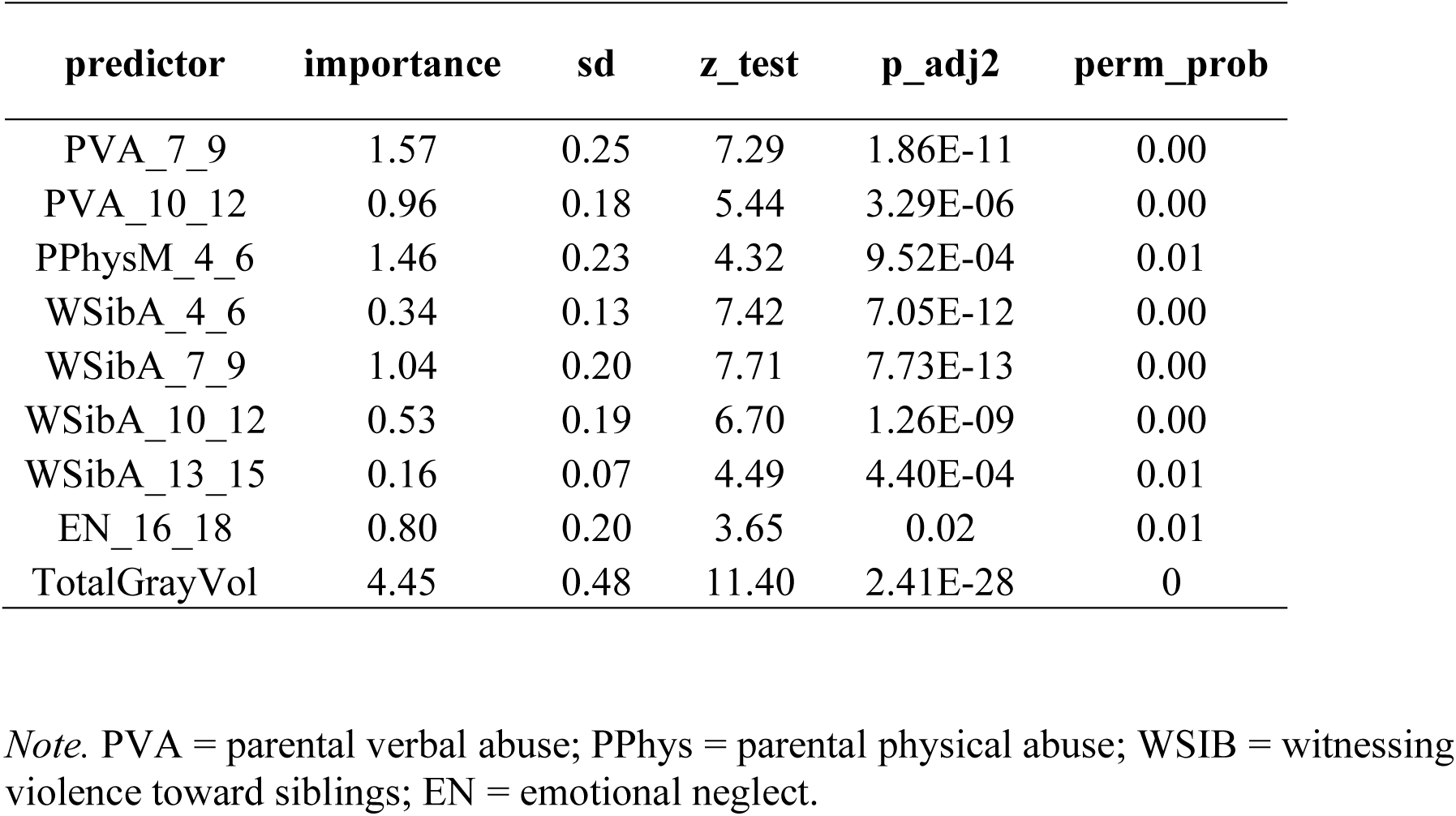
Significant Factors Impacting Brain Age Acceleration.

We then used the results of the random forest models to determine the directionality of effects at each sensitive time point for each type of maltreatment (Fig. 3). Greater exposure to each maltreatment subtype was associated with greater acceleration in brain aging at each sensitive time point, with the exception of parental physical abuse at ages 4 to 6 years, which showed a negative association between brain aging and abuse exposure severity. We also calculated incidence rate ratios (IRRs) for each significant maltreatment type/age-range combination and observed that every one-unit increase in any type of maltreatment severity was associated with deviations in brain age from chronological age by a range of 4% to 30% (Table 2).

**Table 2.** Incidence Ratios and Confidence Intervals for Maltreatment Exposure vs. Brain Age Acceleration.

| **Subtype** | **Age Range**<br>**(years)** | **Incidence**<br>**Rate Ratio**<br>**(95% CI)** | **CI_lower** | **CI_upper** |
| --- | --- | --- | --- | --- |
| PPhys | 4 - 6 | 0.96 | 0.9 | 1.03 |
| PVA | 7-9 | 1.11 | 1.02 | 1.2 |
| PVA | 10-12 | 1.11 | 1.03 | 1.2 |
| EN | 16-18 | 1.09 | 1.01 | 1.17 |
| WSibA | 4-6 | 1.3 | 1.12 | 1.53 |
| WSibA | 7-9 | 1.2 | 1.08 | 1.35 |
| WSibA | 10-12 | 1.28 | 1.12 | 1.48 |
| WSibA | 13-15 | 1.26 | 1.08 | 1.5 |

*Note.* PVA = parental verbal abuse; PPhys = parental physical abuse; WSIB = witnessing violence toward siblings; EN = emotional neglect.

**Figure 3.**
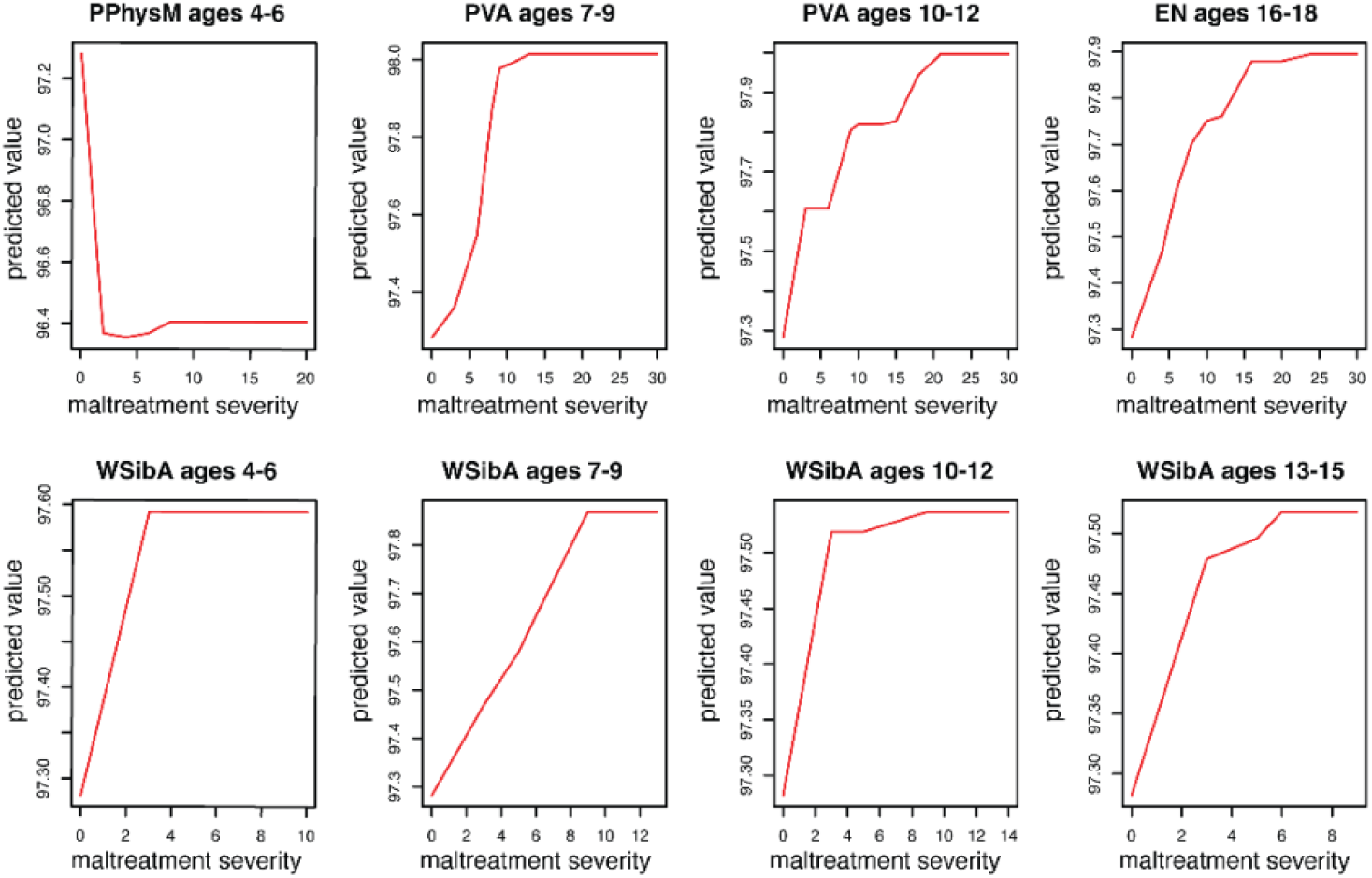
Dose-response effects for sensitive periods of maltreatment. Dose-response plots are shown to depict the directionality of effects for the significant type/times from the random forest regression. Horizontal axes represent severity scores while vertical axes repre­sent scaled, predicted model values. Overall, greater severity was primarily associated with faster brain age acceleration during adulthood. Physical abuse at age 5, however, showed an inverse effect in which greater severity was associated with decelerated brain age. Abbrevia­tions are as follows: PVA, parental verbal abuse; PPhys, parental physical abuse;WSibA, witnessing violence toward siblings; EN emotional neglect;

## DISCUSSION

The goal of the present study was to determine if exposure to childhood maltreatment is associated with accelerated brain aging in adulthood and whether the timing of exposure to various forms of childhood maltreatment relates to differential outcomes in brain aging. In a cohort of adult women, we observed that brain aging was related to the composite severity of maltreatment, consistent with earlier work. Furthermore, we observed sensitive periods for parental physical abuse (4 to 6 years), witnessing sibling violence (4 to 15 years), parental verbal abuse (10 to 12 years), and parental emotional neglect (16 to 18 years). Overall, greater severity of maltreatment was associated with accelerated brain aging, with the exception of physical abuse, which showed a negative association with brain aging from age 4 to 6 years. These results suggest that brain changes, initiated during specific time windows in childhood, induce long-term changes that are evident in adulthood. To the best of our knowledge, such sensitive periods linking childhood maltreatment to brain aging have not been examined previously.

### Sensitive periods for parental verbal abuse

We observed significant associations between parental verbal abuse and accelerated brain aging at 7 to 9 and 10 to 12 years of age. These sensitive periods for brain aging align well with those relating emotional/verbal abuse to other neuropsychiatric outcomes. For example, earlier work showed that emotional abuse experienced between 7 to 12 years of age was a strong predictor of psychological and somatoform dissociation symptoms in PTSD ^44^. Similar effects have also been observed specifically in women during the postpartum period, where parental verbal abuse at age 12 significantly predicted lifetime major depressive disorder ^45^. Work from our own group has also observed significant effects of emotional abuse experienced near this period on depressive symptoms in young adulthood^29^. Here, we extend these findings by demonstrating a link between emotional/verbal abuse experienced during a similar window and adult brain aging. These findings suggest that exposure to emotional/verbal abuse may produce effects that impact multiple, parallel processes related to mental well-being, including both psychiatric symptoms and accelerated brain aging.

### Sensitive periods for the effects of parental physical abuse

We found that exposure to parental physical abuse at ages 4 to 6 years was associated with alteration in brain aging in adulthood. This period is generally parallel to that observed in other work relating physical abuse to psychiatric outcomes. Dunn and colleagues (2013) found that individuals exposed to physical abuse around age 5 years had a 77% higher risk for depression ^46^. Manly and colleagues (2001) also found significant effects for the impact of physical abuse experienced during the preschool age window (3-5 years) on externaling behaviors during later childhood ^47^. There is also evidence linking exposure to physical abuse before age 12 years to increased risk for adult psychotic disorder ^48^. These findings across studies demonstrate the significance of experiences during early developmental periods on adult brain development and psychopathological outcomes.

Interestingly, we found that experiencing physical abuse between ages 4 to 6 years was associated with decreased brain aging. This finding may have multiple explanations. While maltreatment has generally been linked to accelerated pubertal development, the impact on brain maturation appears more complex and non-uniform across the brain ^49–51^. Thus, our results may be in part due to differential impacts on individual brain regions. For example, our finding aligns with earlier work showing delayed maturation of emotional circuitry regions in girls aged 8 to 18 years with histories of physical abuse ^13,15^. However, work focusing specifically on the maturation of emotional circuity in relation to early life adversity is mixed, with some reports indicating accelerated development ^9,52^, while others report delayed development ^13,15,53^. Further, exposure to physical abuse in this age range was associated in early adulthood with a blunted amygdala response to threat, whereas postpubertal exposure to maltreatment was associated with an enhanced response ^30^. Blunted response to threats may exert a beneficial anti-aging effect by reducing threat-associated surges in stress hormones and allostatic load. This suggests a need to disentangle the sources of variability that may contribute to more global associations between maltreatment and brain age, including the contribution of individual regions, timing, and phenotypic variation such as internalizing vs. externalizing psychopathology ^13^.

### Sensitive periods for witnessing violence toward siblings

Witnessing sibling violence showed sensitive periods spanning ages 4 to 15 years. Historically, the study of this form of maltreatment has received limited attention, despite being an important dimension of early life adversity. Limited prior work has demonstrated that witnessing violence towards siblings is associated with ratings of depression, anger-hostility, dissociation, anxiety, somatization, and ‘limbic irritability’ ^54^. The same study found that witnessing violence towards siblings was a stronger predictor of such symptoms than witnessing interparental violence ^54^. These combined results suggest that future work should give more attention to the effects associated with witnessing violence towards siblings.

### Sensitive period for effects of emotional neglect

Emotional neglect experienced during late adolescence (ages 16 to 18 years) was a significant risk factor for accelerated brain aging. Other work examining the effect of emotional neglect on the development of lifetime major depressive disorder also identified sensitive periods during later adolescence ^45^. Thus, emotional neglect may particularly impact the brain as individuals make the transition towards early adulthood, a developmental period that involves multiple, new demands for autonomous decision-making ^55,56^ Frontal regions, in particular, tend to reach maturation during this period ^4^ and are important to decision-making regarding lifestyle, identity, and career choices ^55,56^. Therefore, one question for future work is whether the effects of emotional neglect on brain aging are brain-wide or are more specific to the maturation of frontal regions.

### Limitations and future directions

Study limitations should also be considered, which stand as open opportunities for future directions. One limitation of the current study is its retrospective design, relying on participants to recall maltreatment experiences at specific ages. Such recollections may be susceptible to recall bias, as memories of childhood events, especially those associated with trauma, can be fragmented, reinterpreted, or suppressed over time, although reported events have been remarkably consistent on test-retest. Nevertheless, future studies would benefit from longitudinal designs that may also consider other corroborating reports (such as those of siblings, or other caregivers, and child welfare records). An additional limitation is that this sample consisted of postpartum women under the age of 45. Future studies are needed to assess whether similar results would be obtained in male samples, mothers outside of the postpartum period, non-child-bearing women, or older adults. Lastly, in the current sample, ten types of maltreatment were investigated using the Maltreatment and Abuse Chronology of Exposure scale. Given that we did not pre-screen for the prevalence of each maltreatment subtype in recruitment, some maltreatment subtypes had higher frequency than others (supplemental figure S1). Future work may require more targeted recruitment to include sufficient representation of specific forms of maltreatment.

## Conclusion

The current study contributes to our understanding of the enduring effects of early life adversity and augments our knowledge of how the human brain is shaped by experience.

Importantly, our findings identified age-related periods of particular sensitivity to the effects of several forms of childhood maltreatment on brain aging. Such results need replication, but point to the complexity of maltreatment effects on brain development and underscore the need for nuanced approaches to the study of brain aging to more fully understand underlying mechanisms contributing to the effects observed here. These results also suggest a set of time periods during which screening or clinical attention may help mitigate the long-term risks of childhood maltreatment.

## Declaration of Competing Interests

### Ethical Approval Statement

This study was approved by the Partners Healthcare Institutional Review Board [IRB Protocol #: 2014P002522] and parents provided written informed consent.

## Funding Sources and Acknowledgements

The authors would like to acknowledge support from the following funding resources: National Institute of Child Health and Human Development grants R01HD079484, K00HD111352-03, National Institute of Mental Health K01MH129828, R01MH108665, the Brain Behavior Research Foundation, and a grant from the Mental Wellness Foundation. We would like to thank the families whose interest and participation made this work possible. We would also like to express our appreciation for the hard work and dedication of the study staff who were responsible for subject recruitment and behavioral data acquisition, including Carrie Heldstedt, Rachael Phillips, Molly Rothenberg, Ilana Shiff, Mariya Patwa, Molly Cunningham, Lina Dimitrov, Mallika Rajamani, Danielle Farrell, Sarah Immelt, and Sommer Jaber.

