## Supplemental Materials for "Childhood maltreatment and brain aging during adulthood"

**List of Content:**

**Supplemental Table S1.** Characteristics of Research Participants

**Supplemental Methods 1.** Participant exclusion criteria

**Supplemental Methods 2** Brain age calculation

**Supplemental Methods 3** Random forest regression

**Supplemental Table S2.** Full Random Forest Results

**Supplementary Figure S1.** Percent cases

**This supplementary material has been provided by the authors to give readers additional information about their work.**

**Supplemental Table S1.** Characteristics of Research Participants

| | Finding (n=92) | % (N) |
|---|---|---|
| **Age** | | |
| mean | 32.38 | --- |
| sd | 4.27 | --- |
| **Ethnicity** | | |
| Hispanic | 10 | 10.9% |
| Non-Hispanic | 78 | 84.8% |
| Other | 4 | 4.3% |
| **Race** | | |
| Asian American | 7 | 7.6% |
| Biracial/Multiracial | 8 | 8.7% |
| Black / African American | 6 | 6.5% |
| Caucasian/White | 70 | 76.1% |
| Native Hawaiian/Pacific Islander | 1 | 1.1% |
| **Education** | | |
| Associate degree | 6 | 6.5% |
| Bachelor's Degree | 22 | 23.9% |
| Doctoral Degree | 17 | 18.5% |
| High school graduate | 11 | 12.0% |
| Master's Degree | 34 | 37.0% |
| not a high school graduate | 1 | 1.1% |
| unspecified | 1 | 1.1% |
| **Smoking status** | | |
| non-smoker | 73 | 79.30% |
| smoker | 10 | 10.90% |
| unspecified | 9 | 9.80% |
| **Annual Family Income** | | |
| 0 to $15,000 | 7 | 7.6% |
| $16,000 t0 $25,000 | 7 | 7.6% |
| $26,000 to $50,000 | 10 | 10.9% |
| $51,000 to $75,000 | 12 | 13.0% |
| $76,000 to $100,000 | 16 | 17.4% |
| $101,000 to $150,000 | 26 | 28.3% |
| $151,000 to $200,000 | 11 | 12.0% |
| $201,000 or more | 3 | 3.3% |
| **Maltreatment, MACE score** | | |

| | | |
|---|---|---|
| 0,No exposure | 43 | 46.7% |
| 1, Exposure to 1 types | 23 | 25.0% |
| 2, Exposure to 2 types | 9 | 9.8% |
| ≥3, Exposure to ≥3 types | 17 | 18.5% |
| ≥1, Exposure to ≥1 type | 49 | 53.3% |
| **Total Gray Matter Volume (mm³)** | | |
| mean | 609501.35 | |
| sd | 40623.26 | |

Abbreviation: MACE, Maltreatment and Abuse Chronology of Exposure

### Supplemental Methods

#### *Participant exclusion criteria*

This study utilized data from the Mother-Infant Neurobiological Development (MIND) study. Recruitment was conducted via community flyers, prenatal classes, and local birth records and participants were stratified using the Adverse Childhood Experiences questionnaire [1] such that approximately half of the sample had experienced one or more forms of childhood maltreatment. The key criteria for the MIND study were: ages 18 to 44 years, with English as their primary language, and having given birth to healthy newborns. Exclusion criteria for the main study were: a) English not a primary language spoken at home, b) maternal age over 44 years at the time of infant birth, c) infant born before 36 weeks gestation and/or weighing less than 2500 g, and d) infant congenital disorder/condition. Mothers with a history of head trauma or loss of consciousness lasting more than 20 minutes, those with concussions and standard MRI exclusions (e.g. – metal implant), were not included in the neuroimaging protocol. Participants were also required to abstain from alcohol, drugs, or medications (except contraceptives) for at least two weeks prior to scanning and to have had no history of psychotropic medication use before age 18.

#### *Brain age calculation*

Brain predicted age was calculated using a previously published model through the brainageR package (version 2.1) [2,3]. The brainageR package can be applied to raw, unprocessed T1-weighted MRI images and is publicly available at: https://github.com/james-cole/brainageR. Processing was performed using a combination of software packages, including R version 4.3.1, MATLAB version 2023a, SPM12, and FSL 6.0.6.5. 8. Following pre-processing that included normalization and segmentation of gray matter, white matter, and cerebrospinal fluid, the brainageR package was used to generate brain-predicted age for each participant.

*Random forest regression*

Random forest regression generates a set of random decision trees, each built from a distinct subset of the data and restricted by the number of predictors that can be evaluated at each decision node. The importance of each variable is determined by examining how the fit of the model degrades after its permutation. Under this framework, more important variables produce a more substantial decrease in overall model fit when permuted, whereas less important variables produce very little change in the overall fit when permuted. Here, we implemented random forest regression analysis using in-house code written in *R* (*version 4.3.1*) based on the *cforest* function from the *party* R package, version 1.3.13 http://party.r-forge.rproject.org/). A detailed summary of model results is provided in the tables below.

**Supplemental Table S2**

Full Random Forest Results

| predictor | importance | sd | z_test | p_adj2_BH | perm_prob | lasso | lasso_ridge_p4 |
|---|---|---|---|---|---|---|---|
| E_Dep_1_3 | 0.08621538 | 0.13508211 | 0.06622392 | 0.9850009 | 0.32 | 0 | 0.21097864 |
| E_Dep_4_6 | 0.04600868 | 0.12140331 | -0.0205694 | 0.9850009 | 0.38 | 0 | 0 |
| E_Dep_7_9 | 0.42847158 | 0.14052815 | 1.30047594 | 0.41450978 | 0.09 | 0 | -0.1318668 |
| E_Dep_10_12 | 0.67453653 | 0.21690638 | 1.60264434 | 0.29730872 | 0.068 | 0 | -0.1408872 |
| E_Dep_13_15 | 0.87253928 | 0.24636602 | 2.37513976 | 0.07518131 | 0.03 | 0 | 0 |
| E_Dep_16_18 | 1.38133369 | 0.25119237 | 4.00568025 | 4.64E-04 | 0.006 | 0.16751281 | 0.58686751 |
| P_Dep_1_3 | 0.01072431 | 0.07819085 | -0.1234554 | 0.9850009 | 0.418 | 0 | 0 |

| | | | | | | | |
|---|---|---|---|---|---|---|---|
| P_Dep_4_6 | 0.05097051 | 0.08405827 | -0.0187997 | 0.9850009 | 0.368 | 0 | -0.1510707 |
| P_Dep_7_9 | 0.09388855 | 0.09927522 | 0.08505346 | 0.9850009 | 0.264 | 0 | 0 |
| P_Dep_10_12 | 0.05773064 | 0.09559002 | -0.0736288 | 0.9850009 | 0.356 | 0 | 0 |
| P_Dep_13_15 | -0.0059559 | 0.08459966 | -0.2545832 | 0.9850009 | 0.476 | 0 | 0 |
| P_Dep_16_18 | -0.0188875 | 0.07644713 | -0.2835258 | 0.9850009 | 0.508 | 0 | 0 |
| E_Threat_1_3 | -0.0125587 | 0.07143297 | -0.1146307 | 0.9850009 | 0.396 | 0 | 0 |
| E_Threat_4_6 | 0.61167242 | 0.18463863 | 1.39808013 | 0.37405155 | 0.072 | 0 | 0.11486274 |
| E_Threat_7_9 | 2.16065262 | 0.26326884 | 4.37117317 | 1.85E-04 | 0.008 | 0.06332995 | 0.16453773 |
| E_Threat_10_12 | 1.24938143 | 0.280054 | 2.21099715 | 0.09012011 | 0.028 | 0.00645701 | 0.05869821 |
| E_Threat_13_15 | 0.86695831 | 0.21976171 | 1.40683642 | 0.37405155 | 0.064 | 0 | 0.03786986 |
| E_Threat_16_18 | 0.5753138 | 0.21547816 | 1.02967629 | 0.53499175 | 0.098 | 0.03304226 | 0.0837767 |
| P_Threat_1_3 | -0.2374239 | 0.0805065 | -0.6175846 | 0.80527376 | 0.866 | 0 | -0.0802686 |
| P_Threat_4_6 | 1.3524676 | 0.22486678 | 2.87661104 | 0.02411824 | 0.018 | 0 | -0.2739874 |
| P_Threat_7_9 | 1.76083692 | 0.21694633 | 4.23379728 | 2.30E-04 | 0.008 | 0 | 0.2053622 |
| P_Threat_10_12 | 0.89956665 | 0.24630555 | 2.2449569 | 0.09012011 | 0.028 | 0.11466893 | 0.24158324 |
| P_Threat_13_15 | 0.31885432 | 0.17042623 | 0.85890955 | 0.6506507 | 0.122 | 0 | -0.0204468 |
| P_Threat_16_18 | -0.128945 | 0.12661468 | -0.5665667 | 0.81572662 | 0.776 | 0 | -0.2328071 |
| income_cat | 0.75710526 | 0.19797143 | 1.20614227 | 0.43533666 | 0.086 | 0 | 0.23105806 |
| TotalGrayVol | 7.45059796 | 0.56442796 | 10.3406717 | 1.38E-23 | 0 | -5.68E-05 | -7.21E-05 |
| BQ_CigarettesAny | 0.82750849 | 0.16952035 | 2.71282487 | 0.03335618 | 0.024 | 4.82422422 | 5.31540831 |
| extr.dur | 0.37288752 | 0.18210776 | 0.78095864 | 0.68656866 | 0.142 | 0 | -0.3482311 |
| extr.mace_mult | 0.71356669 | 0.22383437 | 2.08247945 | 0.1118961 | 0.04 | 0 | 0 |
| extr.mace_sum | 0.55717745 | 0.24379538 | 1.19476323 | 0.43533666 | 0.086 | 0 | -0.2131248 |

**Supplementary Figure S1. Percent cases**

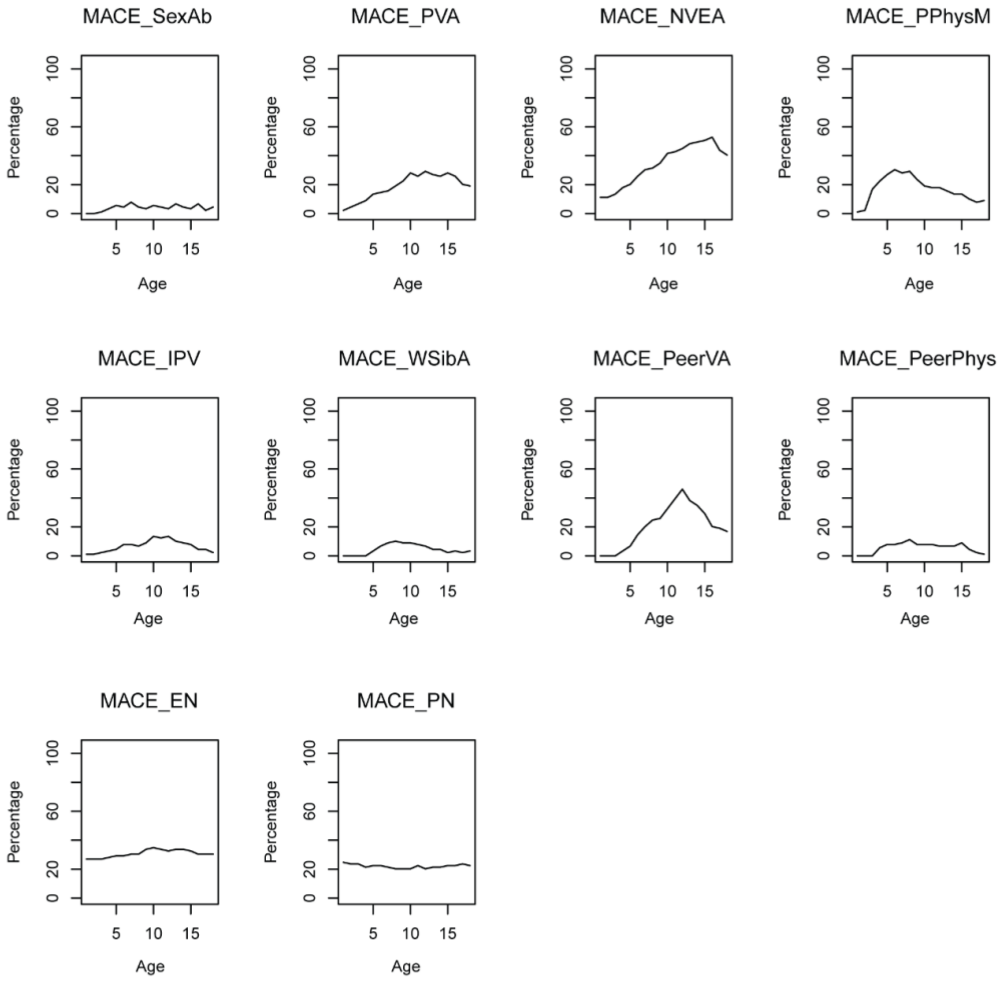


Supplementary Figure S1 - Percent cases across ages. Given the limitations of the random forest regression model, an additional measure was taken to assess how many individuals had history of each type of maltreatment at each timepoint. This is shown above as percentages out of the total number of participants.
